# GABA-A signaling maintains melanocyte stem cell quiescence in larval zebrafish

**DOI:** 10.1101/619056

**Authors:** James R. Allen, James B. Skeath, Stephen L. Johnson

## Abstract

Adult stem cells (ASCs) contribute to long-term homeostasis and regeneration of many adult tissues. Some ASCs proliferate continuously, others remain quiescent awaiting activation. To identify pathways that regulate ASC quiescence and tissue homeostasis, we study melanocyte stem cells (MSCs) that drive vertebrate pigmentation. In larval zebrafish, MSCs are quiescent, but can be recruited to regenerate the larval pigment pattern following melanocyte ablation. Through pharmacological experiments, we found that inhibition of GABA-A receptor function, specifically the GABA-A rho subtype, induces excessive melanocyte production in larval zebrafish. Conversely, pharmacological activation of GABA-A inhibited melanocyte regeneration. We used CRISPR to generate two mutant alleles of *gabrr1*, a subtype of GABA-A. Both alleles exhibited robust melanocyte overproduction, while conditional overexpression of *gabrr1* inhibited larval melanocyte regeneration. Our data suggest that *gabrr1* signaling is necessary and sufficient to maintain MSC quiescence and prevent excessive pigmentation of the larval zebrafish.

## Introduction

Vertebrate animals often rely on undifferentiated precursors to regulate growth and homeostasis of specific tissues. These precursors, adult stem cells (ASCs), undergo long-term self-renewal throughout the lifetime of the organism to maintain the growth and regenerative potential of their target tissue. ASCs are found in many tissues including blood (Bertrand, Kim et al. 2007), muscle (Cheung and Rando 2013), skin (Nishimura, Jordan et al. 2002), and nervous system (Ma, Bonaguidi et al. 2009). While some ASCs continuously proliferate to maintain their target tissue, others remain quiescent or dormant and must be recruited in order to enter a proliferative state, often induced by depletion of differentiated cells in their respective tissues (Li and Bhatia 2011). Understanding the pathways that maintain ASC quiescence and that recruit quiescent ASCs to proliferate is critical to elucidate vertebrate tissue growth and homeostasis.

Zebrafish pigmentation, specifically melanocyte development, is an excellent model system to dissect the genetic and molecular basis of ASC quiescence and recruitment. Both adult melanocytes (Johnson, Africa et al. 1995) and melanocytes that regenerate (Rawls and Johnson 2000) appear to derive from recruitable melanocyte stem cells (MSCs). For example, genetic studies indicate that the embryonic melanocyte pattern develops from direct-developing melanocytes (Hultman, Budi et al. 2009) and is complete by 3 days post fertilization (dpf). Under normal conditions, few new melanocytes develop from 3 dpf until the onset of metamorphosis at approximately 15 dpf (Hultman and Johnson 2010). However, when embryonic melanocytes are removed via chemical (Yang and Johnson 2006) or laser (Yang, Sengelmann et al. 2004) treatment during this time, a rapid and complete regeneration of the melanocyte pattern occurs through activation of cell division in melanocyte precursors, MSCs (Yang and Johnson 2006). MSCs normally lie dormant during larval zebrafish pigmentation, but can be recruited upon loss of differentiated melanocytes. The pathways that regulate MSC quiescence and recruitment are poorly understood.

Forward genetic studies have helped clarify the genetic regulatory hierarchy that controls melanocyte production and MSC proliferation in zebrafish. These studies highlight the importance of three genes in zebrafish pigmentation – the receptor tyrosine kinase *erbb3b*, the transcription factor *mitfa*, and the receptor tyrosine kinase *kita*. Adult zebrafish mutant for *erbb3b*, named *picasso*, exhibit defective melanocyte stripe formation, even though larval *erbb3b* mutants exhibit a wild-type pigment pattern (Budi, Patterson et al. 2008). Critically, when *picasso* mutant zebrafish are challenged for melanocyte regeneration during larval stages, melanocyte regeneration is completely abrogated (Hultman, Budi et al. 2009), suggesting that regenerating melanocytes require *erbb3b* function, while early embryonic melanocytes do not. This finding led to a model wherein a subset migratory neural crest cells directly differentiate into embryonic melanocytes (direct-developing melanocytes), and other *erbb3b*-dependent neural crest cells establish undifferentiated melanocyte precursors, MSCs, that persist throughout zebrafish adult life and can be recruited to form new (stem-cell derived) melanocytes throughout larval and adult stages.

Additional insight into MSCs arose from experiments using a temp-sensitive mutation of the microphthalmia-associated transcription factor (*mitf;* also referred to as melanocyte inducing transcription factor), which is required for all melanocyte development and survival across vertebrate biology (Lister, Robertson et al. 1999, Levy, Khaled et al. 2006). These studies revealed that *mitfa* function was required for the embryonic pigment pattern, but was not required for the survival of MSCs (Johnson, Nguyen et al. 2011). Therefore, while required for melanocyte survival, *mitfa* function is not required for the survival of MSCs that can regenerate larval melanocytes and produces the adult pigment pattern.

The receptor tyrosine kinase *kit* plays key roles during zebrafish pigment patterning. Removal of *kita* function, as seen in the sparse mutant, results in a roughly 50% loss of larval melanocytes, but the adult melanocyte pattern is largely normal (Parichy, Rawls et al. 1999). Thus, *kita* function is required for the development of direct developing melanocytes. *kita* does, however, regulate MSC function. For example, *kita* function is required for melanocyte regeneration during larval stages (O’Reilly-Pol and Johnson 2013) and for melanocyte regeneration in the caudal fin at all stages (Rawls and Johnson 2001, Rawls and Johnson 2003).

GABA is a major inhibitory neurotransmitter that transduces its signal by binding to and activating GABA receptors, such as the GABA-A receptor class (Bormann 2000). GABA-A receptors are voltage-gated chloride channels. When activated, they allow Cl^−^ ions to move down their electrochemical gradient into the cell, which hyperpolarizes the cell and inhibits action potential propagation along axons (Sigel and Steinmann 2012). Although GABA is best known to function as a neurotransmitter, prior studies indicated that GABA can inhibit the proliferation of murine embryonic stem cells (Teng, Tang et al. 2013) and peripheral neural crest stem cells (Young and Bordey 2009). A role for GABA in regulating vertebrate pigment patterning, however, has not been shown.

Here, we show that pharmacological and genetic inhibition of GABA-A receptor function leads to excessive melanocyte production during larval zebrafish development, with the newly produced melanocytes likely arising from MSCs. Conversely, we show that pharmacological or genetic activation of GABA-A signaling inhibits melanocyte regeneration. Our work shows that GABA-mediated signaling promotes MSC quiescence during zebrafish development and highlights the importance of membrane potential and bioelectric sensing in regulating pigment patterning in vertebrates.

## Methods

### Zebrafish Stocks and husbandry

Adult fish were raised and maintained at 14L:10D light-to-dark cycle according to previously standardized protocols (Westerfield 2000). To facilitate melanocyte quantification, the *mlpha* genotype (Sheets, Ransom et al. 2007) was used as wild-type and all experiments were performed in a *mlpha* genetic background unless otherwise indicated. To perform our melanocyte differentiation assay, we used *mlpha* fish carrying Tg(fTyrp1:GFP)^j900^ (Tryon and Johnson 2012). To genetically ablate melanocytes, we used *mlpha* fish homozygous for the temperature-sensitive *mitfa^vc7^* mutation (Johnson, Nguyen et al. 2011). The *kit*^b5^ allele (Parichy, Rawls et al. 1999) in a *mlpha* background was used in the study, and we used the *mlpha* background to generate our two CRISPR-based mutations in *gabrr1*. Embryos of each genotype used in the present study were generated from *in vitro* fertilization.

### Pharmacological Reagents and drug screening

Drugs used in the study were purchased from commercial vendors (See Table S1). Each drug was handled according to manufacture guidelines, but in general each compound was dissolved in a solvent (dimethyl sulfoxide: DMSO or water) to a stock concentration of 20 mM (if possible). For drug experiments, 10-12 embryos were placed into 24-well plates with approximately 2 ml Egg water (60 mg / L Instant Ocean: 0.06 ppt). The stock solution of each drug was then diluted and added to 2 ml Egg water to the indicated concentration of the experiment to a final vehicle concentration of 0.5% DMSO or water (See Table S1). For our melanocyte differentiation assay, we used phenylthiocarbamide (PTU) at a final concentration of 200 µM in egg water. Experiments were performed as parallel duplicates of each drug treatment.

### Melanocyte Counting

We focused our analysis of melanocyte development on the larval dorsal stripe. We quantified melanocytes along the dorsum, beginning with the melanocytes located immediately caudal to the otic vesicle and along the stripe to the posterior edge of the pectoral fin. Each larvae was analyzed once as a biological replicant for the indicated experiment, and no further analysis on individual fish as technical replicants were used in this study. Larvae with severe morphological defects, such as cardiac edema, were excluded from analysis in the present study. P-values were calculated with student’s T-test in Microsoft Excel to compare statistical significance between each experimental and control group of larvae.

### Generation of CRISPR/Cas9-mediated *gabrr1* mutations

We used a previously described software tool (E-CRISP: http://www.e-crisp.org/E-CRISP/(Heigwer, Kerr et al. 2014)) to design guide RNAs that target the *gabrr1* locus. We chose the target sequence (ggatgaaggagcgcttggag), since it targeted a highly conserved stretch of residues within the *gabrr1* coding sequence (Wang, Hackam et al. 1995). Briefly, we cloned our *gabrr1* target sequence into pT7-gRNA (primer: aattaatacgactcactataggatgaaggagcgcttggaggttttagagctagaaatagc). We used the mMessage mMachine T7 RNA synthesis kit to synthesize non-capped guide RNA targeting *gabrr1*. We used the mMessage mMachine SP6 RNA synthesis kit to generate capped Cas9 mRNA from pCS2-nCas9n (Jao, Wente et al. 2013). We injected one-to-two cell stage embryos with *gabrr1* guide RNA, Cas9 mRNA, and phenol red. Injected embryos were reared to adulthood, and sperm samples were screened for Cas9-induced germline mutations. By amplifying a 500 bp amplicon flanking the *gabrr1* target locus, (tggacgggattaaactgagc; aaaatgcaagacccggagat), digesting with T7 endonuclease I and Hpy166II, then analyzing with gel electrophoresis. T7-digested F_0_ founders with putative *gabrr1* lesions were outcrossed to *mlpha* and reared to adulthood. F_1_ individuals were genotyped to identify mutation, outcrossed to Tg(fTyrp1:GFP)^j900^, and analyzed for PTU melanocyte differentiation.

### Generation of hspl:*gabrr1* transgenic line

We generated a stable transgenic line to conditionally overexpress *gabrr1* under the heat shock promoter Tg(hspl:*gabrr1*). We used PCR-based methods to clone the full-length *gabrr1* cDNA obtained from (zv9 assembly; primers: ggcgatcgcttaattaatgttgagggaaagacagctcca; cctgcaggttaattaatcactgtgagtagatggaccagt) into the Pac1 site in pT2-hsp70l. We used the mMessage mMachine SP6 RNA synthesis kit to synthesize capped transposase mRNA from pCS-TP (Kawakami, Takeda et al. 2004). To create a germline integrated hsp:*gabrr1*, we injected embryos at the one or two cell stage with plasmid, tranposase, and phenol red. Injected F_0_ animals were screened at 2-3 dpf for the clonal marker Ef1:GFP, indicating genomic integration of the construct. GFP^+^ embryos were reared to adulthood, outcrossed to *mlpha*, and the resulting progeny were screened for germline transmission of the xenopus EF1*α*:GFP clonal marker (Johnson and Krieg 1995). We established one stable hspl:*gabrr1* transgenic line.

### Heat Shock Induction

Adult zebrafish containing hspl:*gabrr1* were outcrossed to *mlpha* strains to generate clutches of hspl:*gabrr1*; *mlpha*. From 1-3 dpf, embryos were then treated with the melanocyte pro-drug 4-hydroxyanisole (4-HA: 10 mg/ml in DMSO) to ablate melanocytes. At 3 dpf, 4-HA was washed away, embryos were placed in 50 ml conical tubes, and heat shocked at 37 °C in a water bath for 30 minutes. The heat shock treatment was repeated every 24 hours at 3, 4, and 5 dpf. At 6 dpf, the experiment was terminated and larvae were fixed in 3.7% formaldehyde for melanocyte quantification.

## Results

### GABA-A antagonists increase melanocyte production in larval zebrafish

We sought to explore the molecular regulation of melanocyte stem cell quiescence by searching for drugs that result in excess melanocyte development in the larval zebrafish. Previously, our lab found that the larval pigment pattern develops from direct-developing melanocytes and is largely complete by 3 dpf, but that melanocytes that develop after 3 dpf (Hultman and Johnson 2010) or those that regenerate the pigment pattern following melanocyte ablation (Hultman, Budi et al. 2009) develop from MSCs. We took advantage of this finding and designed a small molecule screen to identify compounds that increase melanocyte output after 3 dpf. Our screen used larvae expressing the melanocyte marker f*Tyrp1:GFP*^j900^, incubated them in solution containing the screened compound and the melanin inhibiting drug PTU, and quantified newly generated melanocytes, which can be uniquely identified based on the lack of melanin (mel^−^) and expression of GFP (GFP^+^), the mel^−^, GFP^+^ melanocytes. We focused on the dorsal larval stripe because we previously found that less than two new melanocytes develop within this region between 3 dpf and 6 dpf (Hultman and Johnson 2010). This infrequent development of new melanocytes provided a low background that allowed us to screen for compounds that induced even a small increase in melanocyte production.

We screened over 500 compounds from the Pfizer repurposing panel and identified a GABA-A receptor antagonist (CP-615003-27) that increased melanocyte production between 3 and 6 dpf. Consistent with previous findings from our lab, zebrafish treated with a vehicle control developed on average 1.05 mel^−^, GFP^+^ newly-formed melanocytes in the dorsal larval stripe (Fig. 1B, 1F). Larvae treated with the GABA-A antagonist CP-615003-27 developed on average 4.0 newly formed mel^−^, GFP^+^ melanocytes in the same region (Fig. 1C, 1F), a significant increase over vehicle control treated fish (Fig. 1F). To confirm the effect of GABA-A inhibition on melanocyte production and development, we tested two other GABA-A antagonists. Zebrafish treated with the GABA-A antagonist picrotoxin developed on average 3.7 mel^−^, GFP^+^ melanocytes (Fig. 1D, 1F), and zebrafish treated with the GABA-A rho antagonist TPMPA developed on average 4.1 mel^−^, GFP^+^ melanocytes (Fig. 1E, 1F). Our finding that inhibition of GABA-A receptors through distinct GABA-A antagonists increases melanocyte production suggested that GABA-A signaling regulates MSC quiescence in larval zebrafish.

**Figure 1:**
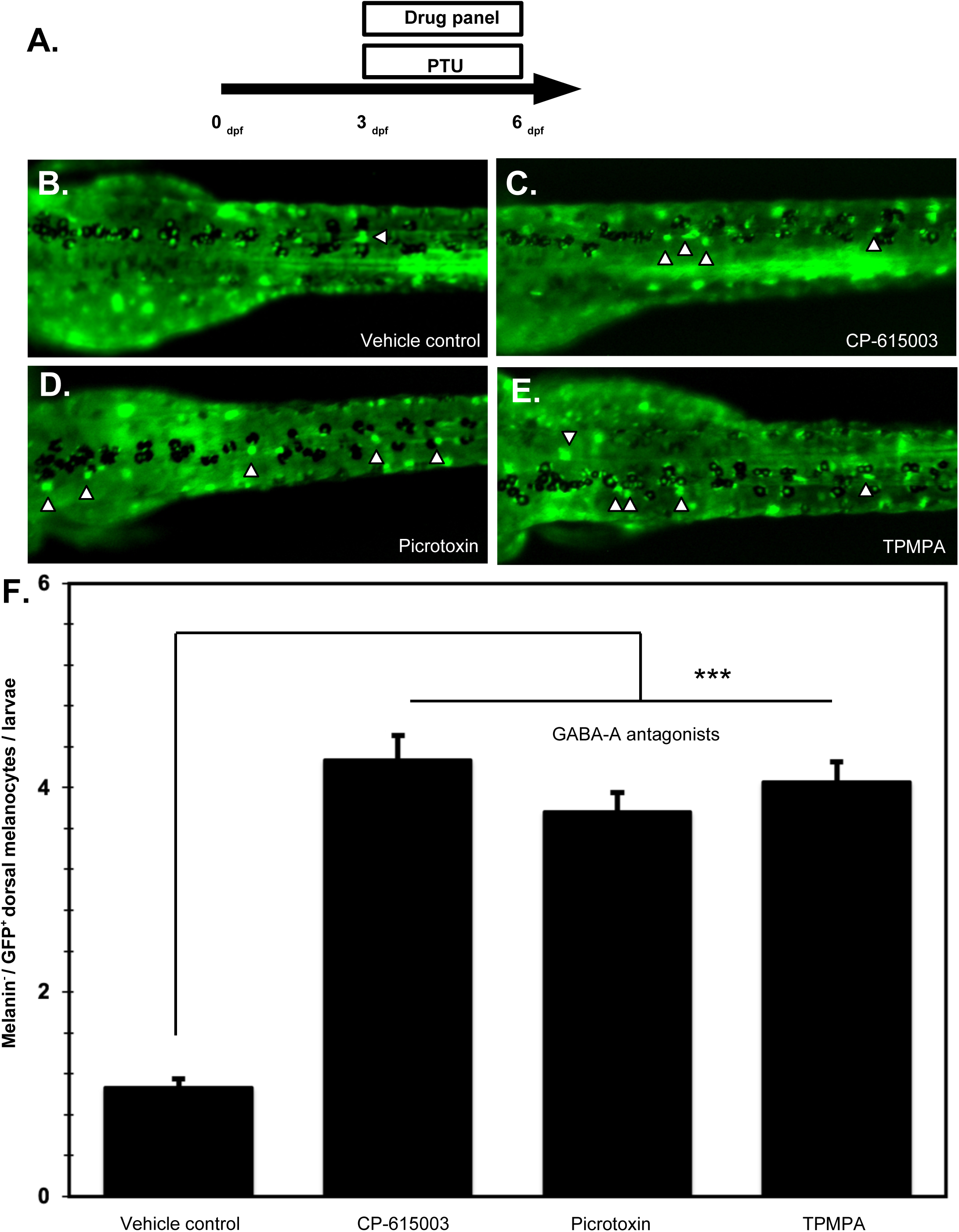
GABA-A antagonists increase melanocyte production in larval zebrafish. (A) Cartoon of experimental timeline for PTU melanocyte differentiation assay. Drugs and PTU are added to zebrafish embryos between 3 – 6 dpf. (B-D) Images of representative 6 dpf larvae treated with vehicle control (B) or GABA-A antagonists CP-615003 (40 µM; C), Picrotoxin (100 µM; D), and TPMPA (100 µM; E). (F) Quantification of the average number of melanin^−^, GFP^+^ dorsal melanocytes for each treatment group. (Vehicle control: 0.92±1.15, S.E.M: 0.08, N=84; CP-615003: 4.27±2.12, S.E.M: 0.23, N=81; Picrotoxin: 3.76±1.38, S.E.M: 0.18, N=55; TPMPA: 4.10±2.14, S.E.M: 0.29, N=52). Error bars represent standard error of mean, each experimental group compared to vehicle control had P-values <0.001 (Students t-test).

### GABA-A antagonist induced melanocytes derive from MSCs

We next asked whether the newly formed melanocytes that develop following GABA-A antagonist treatment arise from a melanocyte stem cell or precursor lineage. Our previous work supported a model that melanocytes within the dorsal stripe primarily develop from undifferentiated MSCs or melanocyte precursors after 3 dpf, suggesting that GABA-A antagonists induce MSCs to produce new melanocytes. However, it remained formally possible that pharmacological inhibition of GABA-A signaling could activate aberrant melanocyte development from a non-stem cell source through an unknown mechanism. To distinguish between these models, we treated zebrafish embryos with either DMSO or the *erbb3* inhibitor AG1478 from 8-48 h.p.f., washed out the drug, and then treated the larvae with solution containing PTU and a GABA-A antagonist from 3-6 dpf (Figure 2A). AG1478-mediated inhibition of *erbb3* activity has been previously shown to inhibit melanocyte regeneration and metamorphic melanocyte development in zebrafish. Early treatment with this small molecule is thought to block establishment of MSCs, removing the developmental source of new melanocytes. Therefore, if new melanocytes arise from MSCs, we predicted that prior AG1478 treatment would inhibit the ability of GABA-A antagonists to induce melanocyte production.

**Figure 2:**
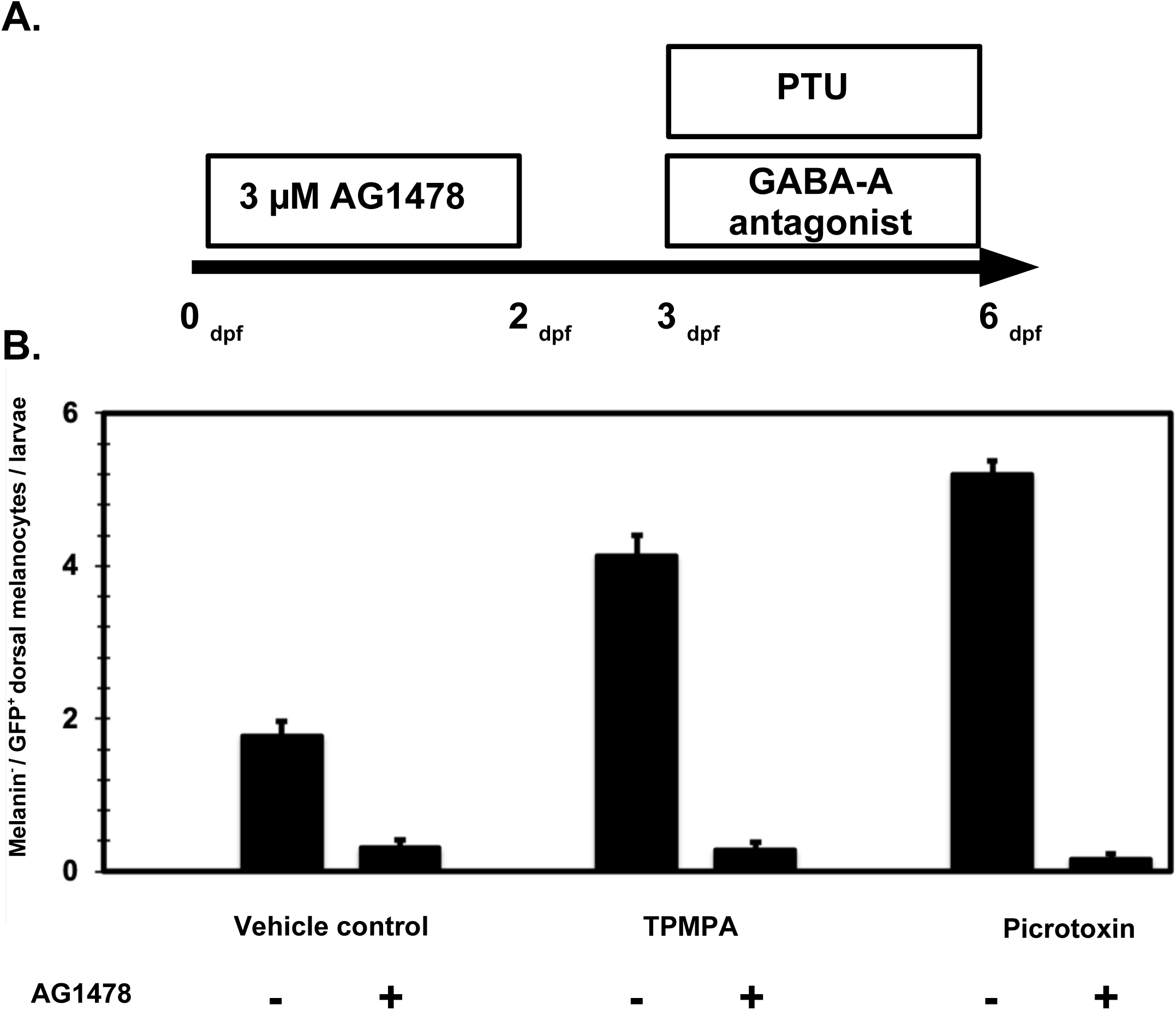
GABA-A antagonist induced melanocytes derive from MSCs. (A) Cartoon of experimental timeline for drug treatment. (B) Quantification of average melanin^−^, GFP^+^ dorsal melanocytes in each group. (Vehicle control: 1.76±1.26, S.E.M: 0.2, N=39; Vehicle control + AG1478: 0.32±0.48, S.E.M: 0.10, N=28; TPMPA: 4.14±1.43, S.E.M: 0.27, N=29; TPMPA + AG1478: 0.28±0.54, S.E.M: 0.11, N=25; Picrotoxin: 5.2±0.99, S.E.M: 0.18, N=30; Picrotoxin + AG1478: 0.17±0.38, S.E.M: 0.06, N=30). Error bars represent standard error of mean, each experimental group compared to vehicle control had P-values <0.001 (Students t-test).

For this analysis, we focused on two representative GABA-A antagonists: TPMPA and Picrotoxin. After each drug treatment, individual larvae were scored for mel^−^, GFP^+^ melanocytes, which we interpret as newly developed melanocytes in the presence of PTU. Zebrafish larvae treated with DMSO and vehicle control developed 1.76 mel^−^, GFP^+^ melanocytes, while larvae treated with AG1478 and vehicle control developed 0.32 mel^−^, GFP^+^ melanocytes (Figure 2B). This result suggested that the AG1478 treatment effectively blocked late (3-6 dpf) melanocyte production. Thus, our PTU assay could detect relatively small changes in melanocyte production, which allowed us to confidently test the combinatorial effects of AG1478 and GABA-A antagonists on melanocyte production. Larvae treated with DMSO and the GABA-A antagonist TPMPA developed 4.1 mel^−^, GFP^+^ melanocytes, but larvae treated with AG1478 and TPMPA developed only 0.28 mel^−^, GFP^+^ melanocytes. Similarly, larvae treated with DMSO and Picrotoxin developed 5.2 mel^−^, GFP^+^ melanocytes, but larvae treated with AG1478 and Picrotoxin developed only 0.17 mel^−^, GFP^+^ melanocytes (Figure 2B). We conclude that GABA-A antagonists induce melanocyte production from *erbb3*-dependent undifferentiated melanocyte precursors.

### Pharmacological activation of GABA-A signaling inhibits melanocyte regeneration

Our data support the model that inhibition of GABA-A receptor signaling increases melanocyte production from undifferentiated precursors. This provided a clear prediction that activation of GABA-A signaling would inhibit melanocyte production. To test this model, we chose to treat larvae homozygous for the temperature sensitive *mitfa^vc7^* allele with drugs that activate GABA-A receptor signaling. When raised at restrictive temperature (32°C), the temperature sensitive nature of the *mitfa^vc7^* allele prevents melanoblast survival, and *mitfa^vc7^* larvae develop no melanocytes. When shifted to permissive temperature (25°C), *mitfa* function is restored and *mitfa^vc7^* larvae exhibit a near complete regeneration of the larval pigment pattern (Johnson, Nguyen et al. 2011). The homozygous *mitfa^vc7^* allele then provided us with temporal control of melanocyte development and regeneration to the test the effects of GABA-A receptor activation.

To determine if GABA-A agonists inhibit melanocyte production, we reared *mitfa^vc7^* larvae to 4 dpf at 32°C, and then down-shifted to 25°C in the presence of a GABA-A receptor drug. Each larvae was scored for the number of dorsal stripe melanocytes present at 7 dpf as a measure of melanocyte regeneration following downshift. *Mitfa^vc7^* larvae treated with vehicle control regenerated 42.2 dorsal melanocytes. However, *mitfa^vc7^* larvae treated with the endogenous ligand GABA or the GABA-A rho agonist GABOB regenerated only 21.2 and 26.8 dorsal melanocytes, respectively (Figure 3B-C; Figure 3G). The reduction of melanocyte regeneration following treatment of GABA-A agonists suggested that direct activation of GABA-A signaling partially inhibited melanocyte regeneration. To further challenge this model, we treated *mitfa*^vc7^ larvae with drugs that indirectly activated GABA-A receptor signaling and challenged for melanocyte regeneration. Larvae treated with the GABA-A partial agonists L-838,417 (Figure 3D; Figure 3G) and MK 0343 (Figure 3F; Figure 3G) regenerated 16 dorsal melanocytes and 18.1 dorsal melanocytes respectively. Similarly, larvae treated with the GABA reuptake inhibitor CI-966, which increases synaptic concentrations of GABA (Ebert and Krnjevic 1990), regenerated only 20.4 dorsal melanocytes (Figure 3E; Figure 3G). The effects of these GABA-A activating drugs were not restricted to the dorsal stripe, and appeared to reduce pigmentation across the ventral and lateral regions of the larvae as well (S. Figure 1). Our data suggest that pharmacological activation of GABA-A receptor signaling inhibits melanocyte production from MSCs.

**Figure 3:**
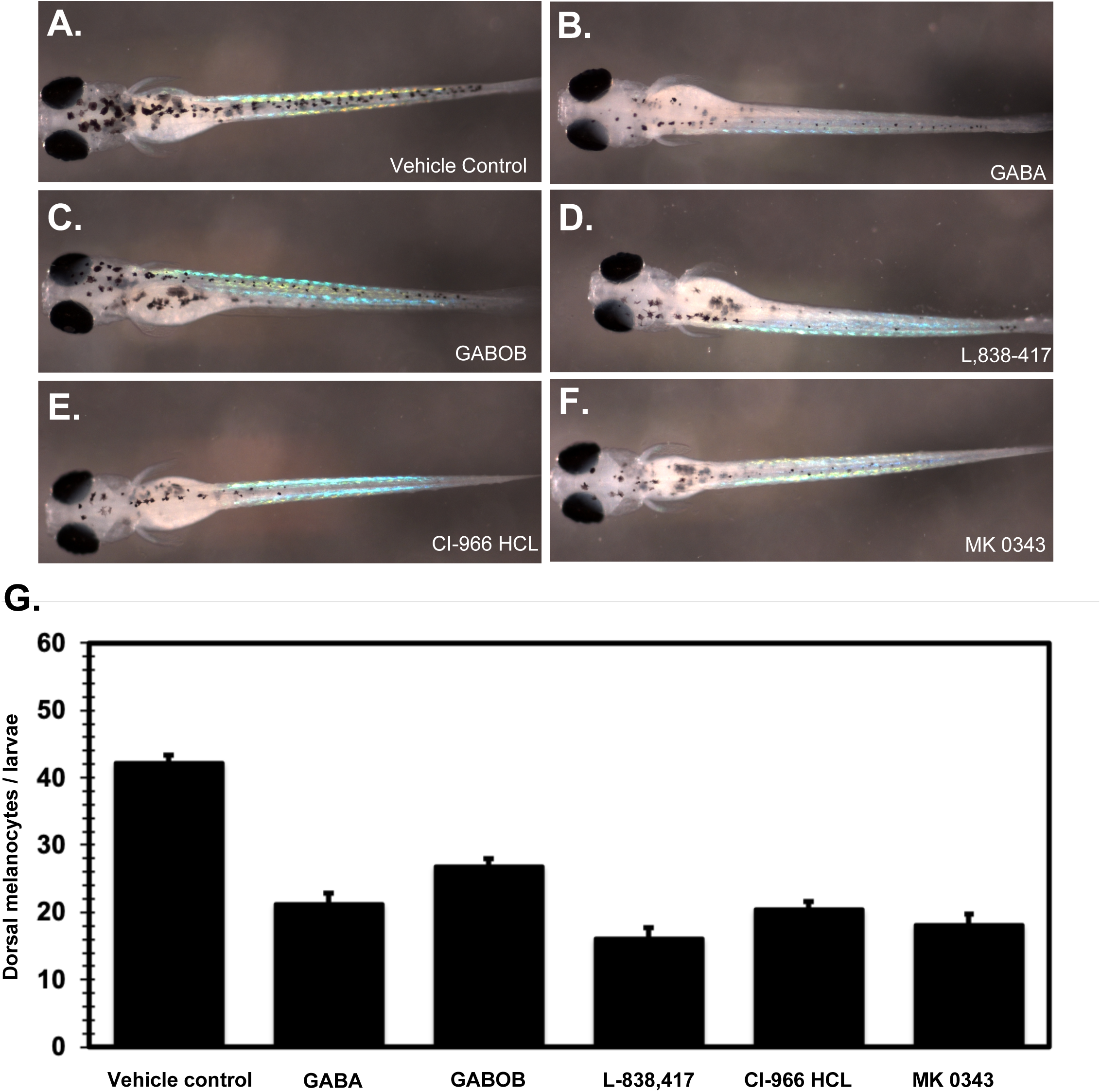
Pharmacological activation of GABA-A signaling inhibits melanocyte regeneration. Images of representative *mitfa*^vc7^ 7 dpf larvae treated with vehicle control (A), GABA (50 mM; B), GABOB (100 µM; C), L, 838-417 (100 µM; D), CI-966 HCL (20 µM; E), and MK 0343 (100 µM; F). (G) Quantification of the average number of dorsal melanocytes in each drug treatment group. (Vehicle control: 42.2±9.38, S.E.M: 1.06, N=78; GABA: 21.2±10.4, S.E.M: 1.61, N=42; GABOB: 26.8±7.71, S.E.M: 1.20, N=41; L,838-417: 16±11.2, S.E.M: 1.73, N=42, CI-966 HCL: 20.4±7.81, S.E.M: 1.19, N=43; MK 0343: 18.1±9.55, S.E.M: 1.61, N=35). Error bars represent standard error of mean, each experimental group compared to vehicle control had P-values <0.001 (Students t-test).

### GABA rho 1 is necessary for restriction of melanocyte production in larval zebrafish

To validate our pharmacology results, we sought to genetically remove GABA-A signaling and assess the impact on melanocyte production. Here, we focused on the GABA-A rho receptor subtype, as the GABA-A rho subtype-specific drug TPMPA yielded a robust increase in melanocyte production. Fortunately, GABA-A rho receptors are homo-pentameric (Martinez-Delgado, Estrada-Mondragon et al. 2010), allowing us to target a single gene to disrupt receptor function. To target GABA-A rho receptor function, we used a CRISPR-based strategy to target the gaba rho 1 (*gabrr1*) gene (see methods). We specifically targeted a region in the ligand-binding domain that is critical for zinc inhibition to increase the likelihood of disrupting endogenous protein function (Wang, Hackam et al. 1995). Using a PCR- and restriction enzyme-based method, we identified two putative *gabrr1* alleles with altered DNA sequence at the targeted site. Sequence analysis of both alleles revealed two *gabrr1* in-frame mutations, both of which delete the conserved residues VHS from position 146-148 of the polypeptide sequence (Figure 4A), with one allele *gabrr1^j247^*, also substituting the lysine at position 149 to glutamic acid.

**Figure 4:**
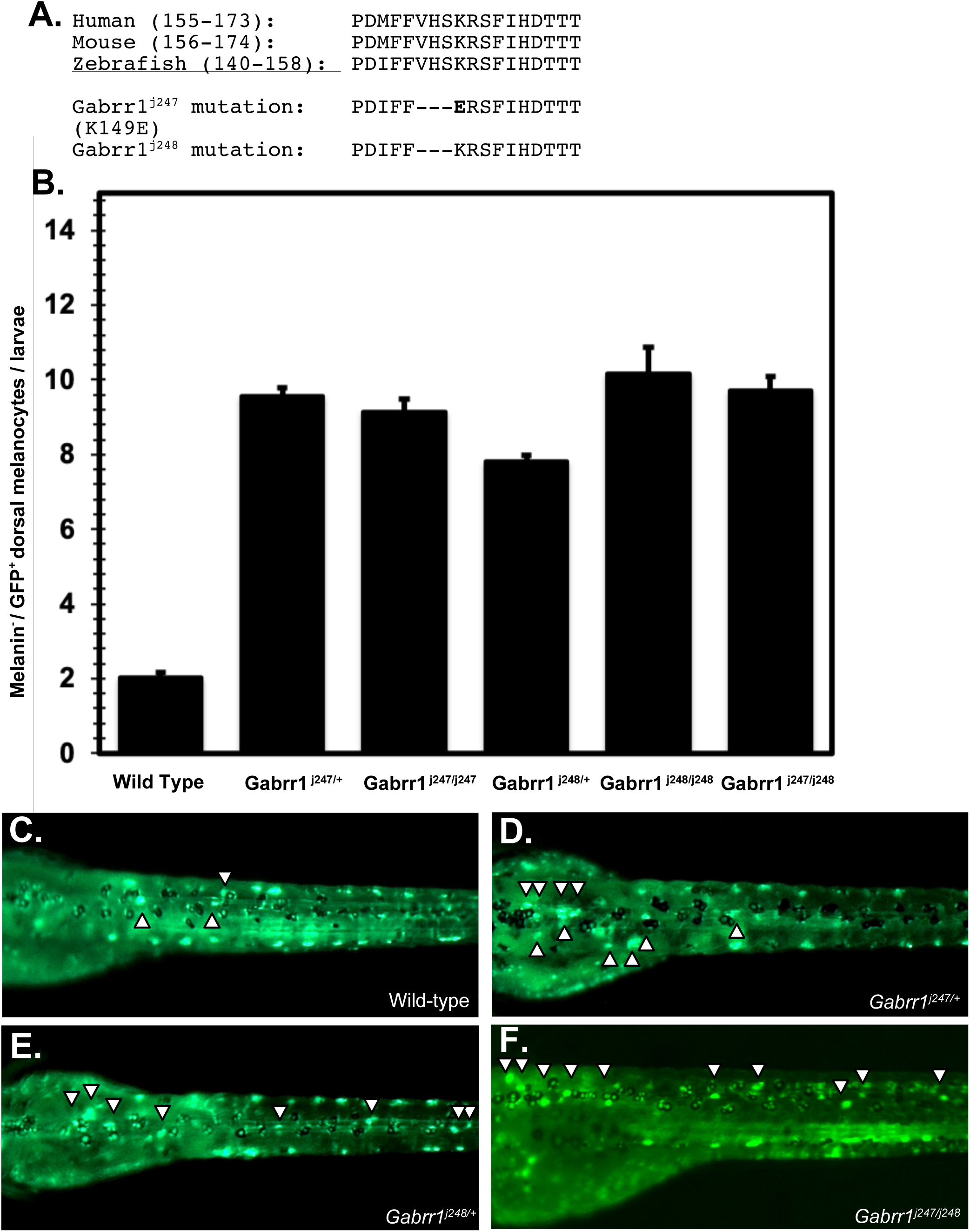
*gabrr1* mutations exhibit a dominant excess melanocyte phenotype during larval stages. (A) Partial alignment of vertebrate *gabrr1* protein sequence homology within the ligand binding domain, with the amino-acid sequence of the two *gabrr1* alleles generated in the study. (B) Quantification of the average number of melanin^−^, GFP^+^ dorsal melanocytes in each treatment group. (Wild Type: 2.04±0.94, S.E.M: 0.13, N=51; *gabrr1^j247/+^*: 9.56±1.48, S.E.M: 0.23, N=39; *gabrr1^j247/j247^*: 9.13±1.41, S.E.M: 0.36, N=15 *gabrr1^j248/+^*: 7.81±1.17, S.E.M: 0.17, N=48; *gabrr1^j248/j248^*: 10.2±2.54, S.E.M: 0.71, N=13; *gabrr1^j247/j248^*: 9.72±1.99, S.E.M: 0.37, N=29). Representative images of 6 dpf wild-type (C), *gabrr1^j247/+^* (D), *gabrr1^j248/+^* (E) and *gabrr1^j247/j248^* (F) larvae. Error bars represent standard error of mean, each experimental group compared to wild type had P-values <0.001 (Students t-test).

To determine if genetic reduction of *gabrr1* function altered melanocyte development, we outcrossed carriers of each *gabrr1* mutation to f*Tyrp1:GFP*^j900^, treated the F_1_ progeny with PTU from 3-6 dpf, and quantified newly generated dorsal melanocytes. Both alleles demonstrated a robust dominant excess melanocyte phenotype.

Zebrafish heterozygous for the *gabrr1^j247^* allele developed 9.56 mel^−^, GFP^+^ dorsal melanocytes (Figure 4B; 4D), a five-fold increase over wild-type siblings (Figure 4B; 4C), and zebrafish heterozygous for the *gabrr1*^j248^ allele developed 7.81 mel^−^, GFP^+^ melanocytes (Figure 4B; 4E). In the PTU assay, the dominant gabrr1 phenotypes were mostly restricted to the dorsal stripe, as we observed little differences in ventral pigmentation (S. Figure 2). In support of both mutations being dominant negative alleles of *gabrr1*, the excess melanocyte phenotype of larval zebrafish homozygous or trans-heterozygous for the alleles was essentially identical to the heterozygous phenotype of each allele (Figure 4B). We infer that *gabrr1* is necessary to inhibit excessive melanocyte production in the larval zebrafish, suggesting that GABA signaling through *gabrr1* is a key regulatory pathway that normally maintains melanocyte stem cell quiescence in larval zebrafish.

### GABA rho 1 is sufficient to inhibit melanocyte regeneration in larval zebrafish

Our observation that *gabrr1* function is necessary to inhibit melanocyte production led us to test whether over-expression of *gabrr1* was sufficient to inhibit melanocyte regeneration. To address this question, we cloned the *gabrr1* cDNA under control of the heat shock promoter Hsp70l within the Tol2 germ-line transformation vector (Suster, Kikuta et al. 2009) and obtained a stable transgenic line: Tg(Hsp70l:*gabrr1*) (see methods). We then treated Tg(Hsp70l:*gabrr1)* and control larvae with the drug 4-HA from 1-3 dpf to ablate melanocytes, washed the drug out, induced heat shock at 37°C, and then quantified melanocyte regeneration at 6 dpf (Figure 5A). Heat shocked wild-type and Non-heat shocked larvae regenerated 49.9 and 51.1 melanocytes respectively, whereas heat shocked Tg(Hsp70l:*gabrr1)* larvae regenerated 32.3 melanocytes, a roughly 40% reduction in melanocyte production (Figure 5B; 5C). Thus, overexpression of *gabrr1* can repress production of melanocytes during periods of regeneration. The over-expression of *gabrr1* also appeared to inhibit melanocyte production both ventrally and laterally, but this effect was not as obvious as the effect of the dorsal stripe (S. Fig 3). Expression of *gabrr1* then appears sufficient to inhibit melanocyte regeneration in larval zebrafish.

**Figure 5:**
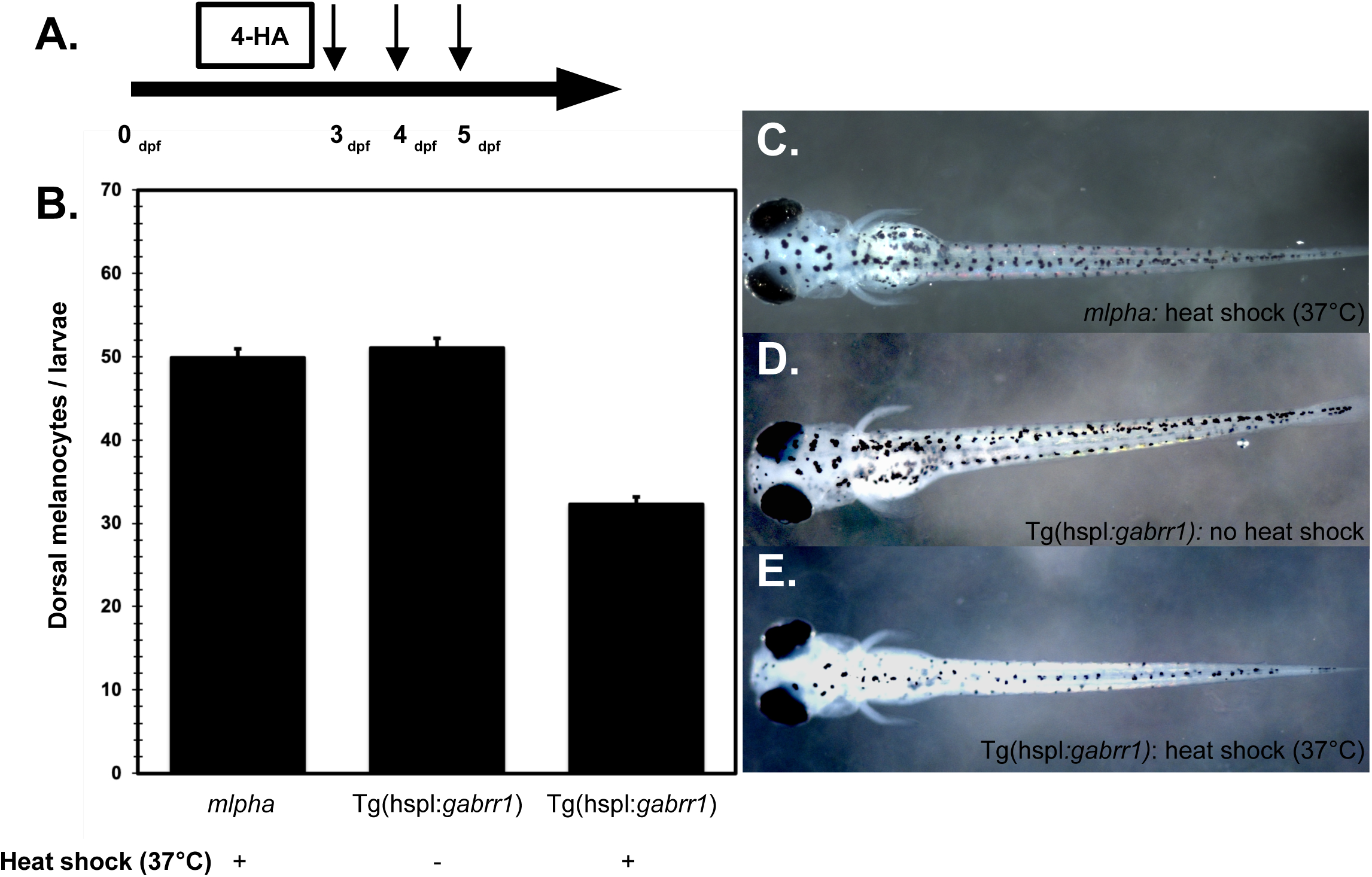
Over-expression of *gabrr1* inhibits melanocyte regeneration in larval zebrafish. (A) Cartoon of experimental timeline. Arrows indicate timing of three 30 minute 37 °C heat shock treatments. (B) Quantification of average dorsal melanocytes in each treatment group. (mlpha + heat shock: 49.9±6.01, S.E.M: 1.06, N=32; Tg(Hsp70l:*gabrr1)*: 51.1±6.86, S.E.M: 1.11, N=38; Tg(Hsp70l:*gabrr1)+* heatshock: 32.3±5.54, S.E.M: 0.81, N=46). Images of representative mlpha + heat shock (C), Tg(Hsp70l:*gabrr1)* (D), and Tg(Hsp70l:*gabrr1) +* heatshock (E) larvae. Error bars represent standard error of mean, Tg(Hsp70l:*gabrr1) +* heat shock compared to mlpha + heat shock had P-values <0.001 (Students t-test).

### Kit signaling and gabrr1

We next determined whether *kita* function was required for GABAergic maintenance of melanocyte stem cell quiescence. Zebrafish heterozygous for the *kita*^b5^ null allele regenerate only about 50% of the larval pigment pattern (O’Reilly-Pol and Johnson 2013), suggesting a reduction of the melanocyte stem cell pool consistent with the effects of *kit* haploinsufficiency observed in mammals (Geissler, Ryan et al. 1988). To test for possible interactions between *kit* and *gabrr1*, we asked whether *kita* haploinsufficiency inhibited the melanocyte over-production phenotype observed in *gabrr1* mutants. We generated control and *kita^b5/+^; gabrr1^j247/+^* double-heterozygous larvae, reared them to 3 dpf, treated them with PTU, and then scored for excess melanocyte production at 6 dpf. As previously observed, wild-type larvae developed 2 excess melanocytes between 3-6 dpf, *kita^b5/+^* larvae developed 1.5 excess melanocytes, and *gabrr1^j247/+^* developed 9 excess melanocytes (Figure 6). Of note, *kita^b5/+^; gabrr1^j247/+^* larvae developed 1.5 excess melanocytes, indicating that the *gabrr1* melanocyte over-production phenotype depends entirely on full *kita* function, suggesting that GABA-A mediated MSC quiescence is restricted within *kita*-dependent melanocyte lineages.

**Figure 6:**
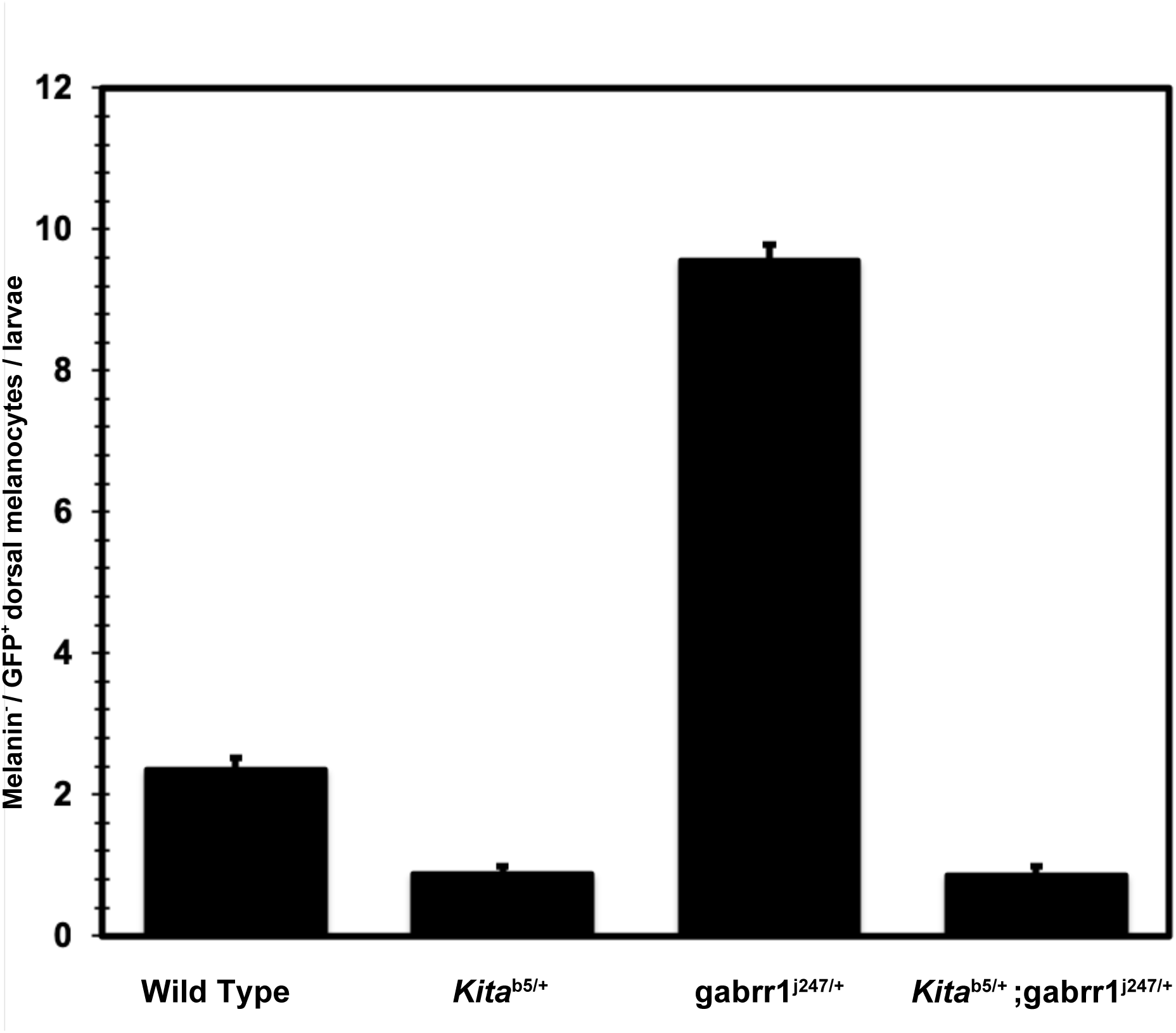
*gabrr1* mediated maintenance of MSC quiescence is sensitive to kit dosage. Quantification of the average number of melanin^−^, GFP^+^ dorsal melanocytes in *gabrr1*^j247/+^ and kit^b5^ heterozygotes. (Wild-type: 2.35±1.03, S.E.M: 0.16, N=42; kit^b5/+^: 0.88±0.81, S.E.M: 0.11, N=56; *Gabrr1*^j247/+^: 9.55±1.43, S.E.M: 0.23, N=40; kit^b5/+^; *gabrr1*^j247/+^: 0.87±0.87, S.E.M: 0.11, N=60). Error bars represent standard error of mean, Each group compared to vehicle control had P-values <0.001 (Students t-test).

## Discussion

Our work provides evidence that GABAergic signaling promotes MSC quiescence in larval zebrafish. Both pharmacological and genetic studies indicate that reduction of GABA-A rho signaling increases melanocyte production, whereas over-expression of GABA-A rho signaling inhibits melanocyte production. Although both classical and recent studies implicate membrane potential in pigmentation and stem cell proliferation, to our knowledge, our study is the first to uncover such a role for GABA-A receptor signaling in vertebrate pigment biology. Below, we propose a model for how GABA-A signaling regulates melanocyte development, discuss the nature of the GABA-A rho mutants, and place our work in the context of old and new studies that highlight the importance of membrane potential and cell excitability in regulating stem cell proliferation and pigmentation.

Our work indicates that GABAergic signaling, directly or indirectly, maintains the MSC in a quiescent state. Prior studies suggest that differentiated larval melanocytes inhibit melanocyte differentiation by suppressing MSC proliferation. For example, chemical or laser ablation of differentiated larval melanocytes induces melanocyte regeneration by promoting the cell division of stem-cell like melanocyte progenitors. In this context, our work provides a conceptual, albeit speculative, model for how the presence of differentiated melanocytes promotes MSC quiescence and how their absence triggers MSC proliferation and melanocyte production. Our pharmacological and genetic data support a model wherein melanocytes release the neurotransmitter GABA, which activates *gabrr1* receptors on the melanocyte stem cell, maintaining the MSC in a quiescent state. Conversely, loss of melanocytes would trigger a reduction in GABA concentration and relieve *gabrr1*-mediated quiescence, triggering MSC proliferation and melanocyte production. Alternatively, GABAergic signaling could act indirectly to regulate MSC quiescence. For example, the melanocyte could release a non-GABA signal, which triggers a GABA to GABA receptor relay in adjacent cells and tissues that ultimately promotes MSC quiescence. Clearly, additional work is required to address whether GABAergic signaling directly or indirectly controls MSC quiescence, but the presence of GABA synthesis enzymes, such as GAD67 mRNA (Ito, Tanaka et al. 2007), in human melanocytes hints that GABA signaling may be an evolutionarily conserved mechanism that regulates vertebrate pigmentation.

### Dominant-negative nature of *gabrr1* alleles

Both of our *gabrr1* alleles exhibit essentially identical melanocyte over-production phenotypes when in the heterozygous or homozygous state. The apparent dominant-negative nature of these alleles was initially surprising. But, both alleles remove a highly conserved triplet of amino acids in the ligand-binding domain of the receptor (Wang, Hackam et al. 1995). Thus, each allele likely produces a non-functional subunit. As GABA-rho 1 receptors function as homo-pentamers, assuming the mutant form of the protein is expressed at roughly wild-type levels, 97% of all *gabrr1* channels would be expected to contain at least one mutant subunit, providing a rational explanation for the dominant nature of the *gabrr1* mutant alleles.

### Do multiple extrinsic pathways regulate MSC quiescence?

When challenged for regeneration, zebrafish larvae produces hundreds of new melanocytes to repopulate the pigment pattern. These new melanocytes derive from a pool of established precursors, MSCs. Though usually quiescent, MSCs are capable of producing hundreds of new melanocytes throughout development. Using clonal analysis, we previously estimated that the developing zebrafish establishes between 150-200 MSCs before 2 dpf (Tryon, Higdon et al. 2011). The MSC pool, though quiescent, then maintains abundance in number and regenerative capability. Complete abrogation of the mechanisms that maintain MSC quiescence would then be expected to generate excess melanocytes proportional to the regenerative capabilities of all MSCs, i.e. generate hundreds of excess melanocytes. Pharmacological or genetic inhibition of GABA-A signaling, however, yields only 8-10 excess melanocytes (Fig. 1; Fig. 4). Thus, the full regenerative capability of larval zebrafish likely involves the concerted actions of multiple pathways that converge on activation of MSC proliferation.

Our genetic studies on *kita* and *gabrr1* suggest GABA-A signaling may regulate a *kita-* dependent pool of MSCs (Tu and Johnson 2010). For example, our prior work indicated that haploinsufficiency for *kita* reduces the available MSC pool by about 50% (O’Reilly-Pol and Johnson 2013). But, haploinsufficiency for *kita* completely suppressed the over-production of melanocytes caused by reduced *gabrr1* function. Our findings suggest that kit signaling is required for all GABA sensitive MSCs, but not all MSCs within the larvae are sensitive to either *kita* or GABA-A signaling. Interestingly, our finding that *gabrr1*-specific melanocyte production is inhibited by *kita* haploinsufficiency also supports a phenomena of *kit* signaling restriction observed in clinical melanomas. Acral (primarily in palms of hands and soles of feet) and mucosal (primarily in oral or nasal cavity and other mucosal surfaces) melanomas are unique melanoma subtypes often associated with oncogenic mutations in c-kit (Curtin, Busam et al. 2006), suggesting that ectopic activation of kit signaling promotes or supports spatially restricted presentations of melanoma. Perhaps the requirement of kit signaling within the *gabrr1-*driven melanocyte lineage of zebrafish is indicative of regulatory pathways that suppress melanocyte production in a region-specific manner. While the functional relevance of this observation remains unknown, it’s clear that complex and distinct aberrations in the melanocyte lineage may play unique roles in melanocyte development and pathology.

Which pathways function in parallel with *gabrr1* to suppress MSC proliferation remain unclear. Recent work completed during the course of our study suggest the endothelin receptor Aa (*ednraa*) acts in parallel to *gabrr1* to maintain MSC quiescence. For example, loss of *ednraa* function leads to ectopic melanocyte production (via a MSC intermediate) specifically within the ventral truck of larval zebrafish (Camargo-Sosa, Colanesi et al. 2019), whereas we find that genetic reduction of *gabrr1* function increased MSC-derived melanocyte production in the dorsal stripe. Intriguingly, pharmacological (S. Fig 1) or genetic (S. Fig 3) activation of *gabrr1* signaling appeared to reduce pigmentation in a larval-wide manner, rather than a region specific effect one. Regardless of the underlying reasons behind the region-specific and larval-wide phenotypes between reduction and activation of *gabrr1* signaling, these studies support the idea that distinct genetic pathways maintain MSC quiescence in a region-specific manner throughout zebrafish development. Clearly, additional work is needed to determine whether other pathways act with *gabbr1* and *ednraa* to promote MSC quiescence, but our work hints that GABA-A mediated quiescence may be a hallmark of vertebrate pigment biology. For example, human melanocytes express enzymes such as aldehyde dehydrogenases (*aldh1a1 and aldh9a1)* that can synthesize GABA (Ganesan, Ho et al. 2008). In addition, *gabrr1* expression is reduced in zebrafish melanoma compared to melanocytes (Venkatesan, Vyas et al. 2018), suggesting that down-regulation of GABA-A signaling could support adult melanoma growth, consistent with our findings in larval melanocyte biology. Whether GABA-A mediated MSC quiescence is a hallmark of vertebrate pigment biology, was abandoned at some point in vertebrate evolution, or is a zebrafish restricted phenomenon is of interest moving forward.

### Bioelectric regulation of MSC quiescence and proliferation

Recent and more classical studies reveal that altering the bioelectric properties, or membrane potential, of progenitor or stem cells may be a fundamental, but poorly understood, developmental mechanism that regulates stem cell activity. For example, pharmacological depolarization of glycine gated chloride channels induces severe hyperpigmentation in xenopus larvae via melanocyte over-proliferation and over-production (Blackiston, Adams et al. 2011). In addition, application of GABA and GABA-A agonists, which hyperpolarizes cells by promoting Cl^−^ influx, has been shown to inhibit proliferation of embryonic stem cells (Teng, Tang et al. 2013) and peripheral neural crest stem cells (Young and Bordey 2009) in mice. Our work complements these studies, as we found that inhibition of *gabrr1* promotes melanocyte production, likely through an MSC intermediate. Thus, altering the membrane potential of Cl^−^ appears to play a key role in regulating vertebrate pigmentation and stem cell proliferation.

Recent work in mice and humans underlines the importance of membrane potential and the bioelectric properties of cells in regulating stem cell activity and development. For example, in the mouse neocortex, neural progenitors become increasingly hyperpolarized as they produce their characteristics cell lineages (Vitali, Fievre et al. 2018). Of note, artificially hyperpolarizing neural progenitor cells induced the premature production of late-stage cell types, revealing a functional link between changes in membrane potential and temporal birth order of cells in the neocortex. In humans, developmental defects in cerebral cortex folding and neuronal migration strongly associate with mutations in Na_v_1.3, a voltage-gated sodium channel primarily expressed at developmental stages composed of early non-transmitting neurons (Smith, Kenny et al. 2018). This suggests that the cellular basis for these neurological defects arises not from altered neuronal transmission, but rather from alterations in the development and migration of neural progenitors and neurons. Together, these studies and our work provide strong evidence that changes in the membrane potential of stem and progenitor cells alters cell division patterns and cellular behavior, highlighting the emerging theme of bioelectric regulation of stem and progenitor cells during development. Future work that can systematically assess the effect of membrane potential on the development of stem cells and their progeny in a broad range of tissues is required to reveal the extent to which this phenomenon occurs in vertebrate biology.

## Acknowledgements

We thank Brian Stephens and Sinan Li for fish husbandry during the majority of the study. We are especially grateful to the Washington University School of Medicine Genetics Department and Washington University Zebrafish Facility for providing critical support in completing this research. We thank Rob Tryon and Ryan McAdow for assistance and guidance generating mutant and transgenic lines. We thank Michael Nonet, Cristina Strong, Charles Kaufman, and Douglas Chalker for critical comments on the manuscript. James R. Allen was a Howard Hughes Medical Institute Gilliam Fellow during the course of this study. This work was funded by NIH RO1-GM056988 to S.L.J. J.B.S was supported by NIH RO1-NS036570.

---

**Table S1:** List of GABA-A receptor pharmacology used in study. Each drug was obtained from the indicated manufacturer and handled according to vendor guidelines.

| **Drug** | **Class** | **Manufacture** | **Conc. In experiment** |
| --- | --- | --- | --- |
| CP-615003-27 | GABA-A Antagonist | Pfizer | 100 µM |
| Picrotoxin | GABA-A Antagonist | Tocris | 100 µM |
| TPMPA | GABA-A Antagonist | Tocris | 100 µM |
| GABA | GABA-A Agonist | Sigma-Aldrich | 100 µM |
| GABOB | GABA-A Agonist | Sigma-Aldrich | 100 µM |
| L, 838-417 | GABA-A partial agonist | Tocris | 1 µM |
| MK 0343 | GABA-A partial agonist | Tocris | 30 µM |
| CI-966 HCL | GABA uptake inhibitor | Tocris | 100 µM |

**Supplemental Figure 1:**
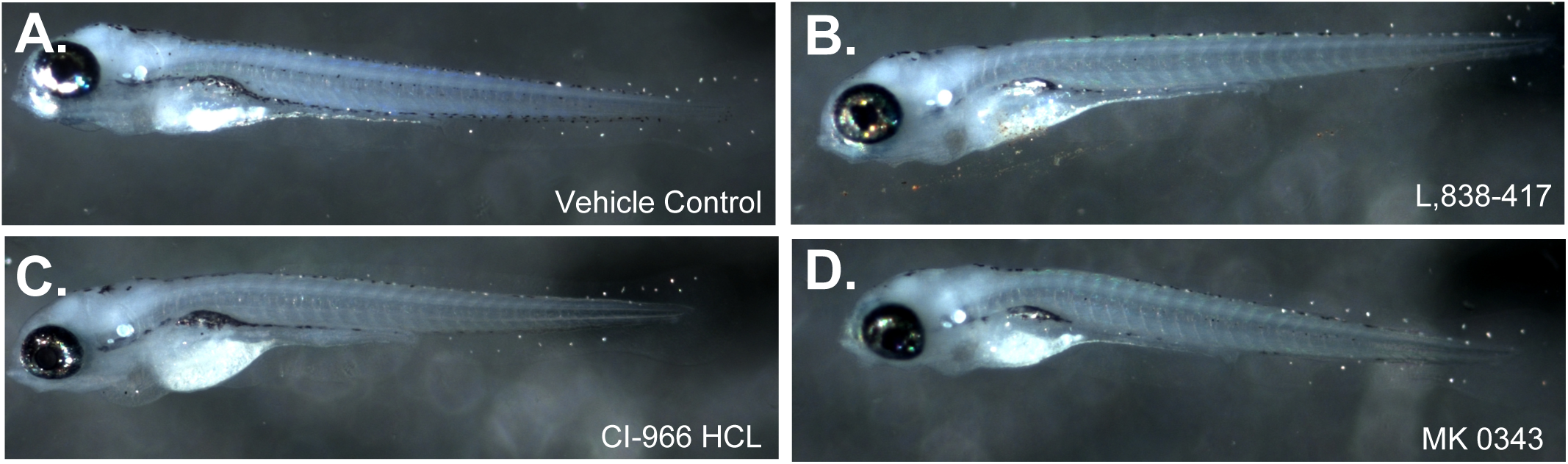
Pharmacological activation of GABA-A reduces larval pigmentation across the body. (A-D) Images of representative *mitfa*^vc7^ 7 dpf larvae treated with vehicle control (A), L,838-417 (B), CI-966 HCL (C), and MK 0343 (D).

**Supplemental Figure 2:**
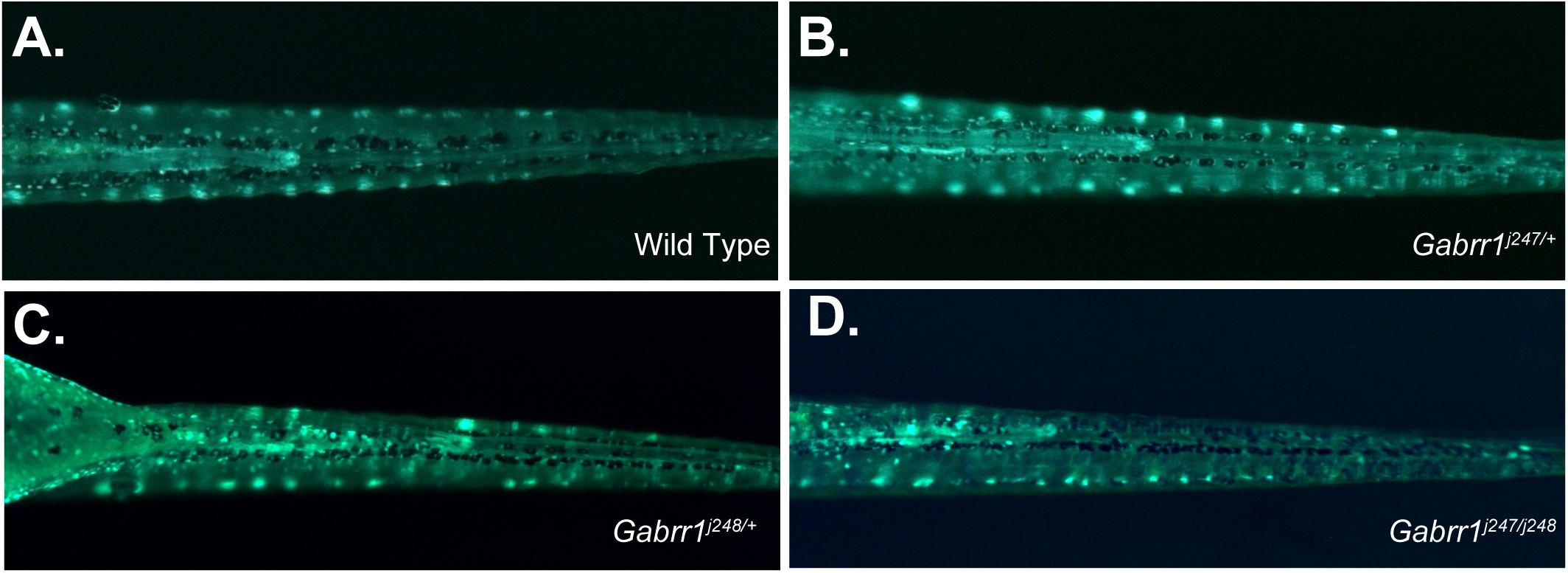
*gabrr1* mutant phenotypes have no visible effect on ventral pigmentation. (A-D) Images of representative 6 dpf wild-type (A), *gabrr1^j247/+^* (B), *gabrr1^j248/+^* (C), and *gabrr1^j247/j248^*(D) larvae.

**Supplemental Figure 3:**
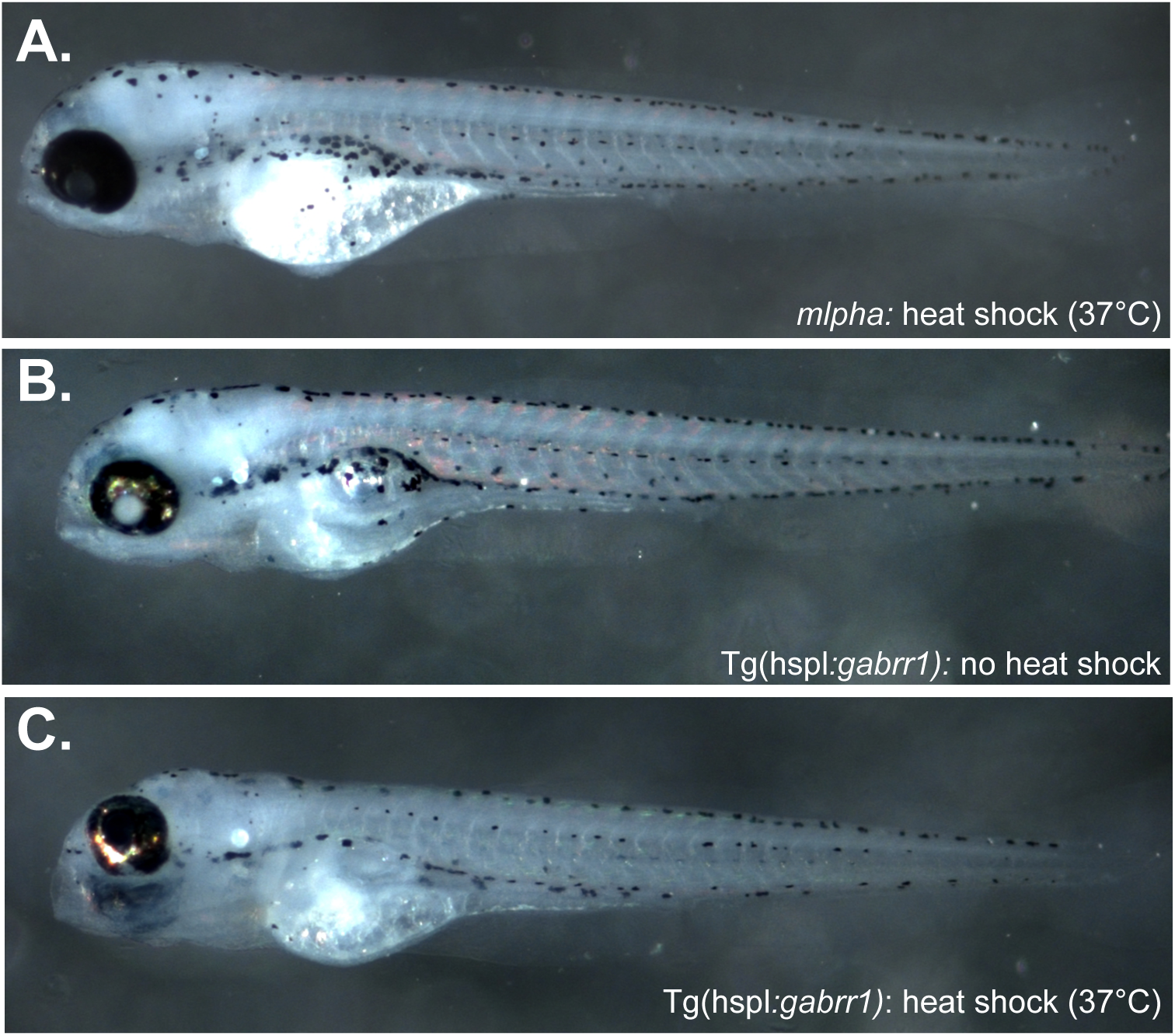
Over-expression of *gabrr1* partially reduces ventral pigmentation. (A-C) Images of representative 6 dpf heat shocked mlpha (A), Tg(Hsp70l:*gabrr1)* (B), and Tg(Hsp70l:*gabrr1)* + heat shock (C).

